# The maternal genetic make-up of the Iberian Peninsula between the Neolithic and the Early Bronze Age

**DOI:** 10.1101/106963

**Authors:** Anna Szécsényi-Nagy, Christina Roth, Guido Brandt, Cristina Rihuete-Herrada, Cristina Tejedor-Rodríguez, Petra Held, íñigo García-Martínez-de-Lagrán, Héctor Arcusa Magallón, Stephanie Zesch, Corina Knipper, Eszter Bánffy, Susanne Friederich, Harald Meller, Primitiva Bueno Ramírez, Rosa Barroso Bermejo, Rodrigo de Balbín Behrmann, Ana M. Herrero-Corral, Raúl Flores Fernández, Carmen Alonso Fernández, Javier Jiménez Echevarria, Laura Rindlisbacher, Camila Oliart, María-Inés Fregeiro, Ignacio Soriano, Oriol Vicente, Rafael Micó, Vicente Lull, Jorge Soler Díaz, Juan Antonio López Padilla, Consuelo Roca de Togores Muñoz, Mauro S. Hernández Pérez, Francisco Javier Jover Maestre, Joaquín Lomba Maurandi, Azucena Avilés Fernández, Katina T. Lillios, Ana Maria Silva, Miguel Magalhães Ramalho, Luiz Miguel Oosterbeek, Claudia Cunha, Anna J. Waterman, Jordi Roig Buxó, Andrés Martínez, Juana Ponce Martínez, Mark Hunt Ortiz, Juan Carlos Mejías-García, Juan Carlos Pecero Espín, Rosario Cruz-Auñón Briones, Tiago Tomé, Eduardo Carmona Ballestero, João Luís Cardoso, Ana Cristina Araújo, Corina Liesau von Lettow-Vorbeck, Concepción Blasco Bosqued, Patricia Ríos Mendoza, Ana Pujante, José I. Royo-Guillén, Marco Aurelio Esquembre Beviá, Victor Manuel Dos Santos Goncalves, Rui Parreira, Elena Morán Hernández, Elena Méndez Izquierdo, Jorge Vega de Miguel, Roberto Menduiña García, Victoria Martínez Calvo, Oscar López Jiménez, Johannes Krause, Sandra L. Pichler, Rafael Garrido-Pena, Michael Kunst, Roberto Risch, Manuel A. Rojo-Guerra, Wolfgang Haak, Kurt W. Alt

## Abstract

Agriculture first reached the Iberian Peninsula around 5700 BCE. However, little is known about the genetic structure and changes of prehistoric populations in different geographic areas of Iberia. In our study, we focused on the maternal genetic makeup of the Neolithic (∼ 5500-3000 BCE), Chalcolithic (∼ 3000-2200 BCE) and Early Bronze Age (∼ 2200-1500 BCE). We report ancient mitochondrial DNA results of 213 individuals (151 HVS-I sequences) from the northeast, central, southeast and southwest regions and thus on the largest archaeogenetic dataset from the Peninsula to date. Similar to other parts of Europe, we observe a discontinuity between hunter-gatherers and the first farmers of the Neolithic. During the subsequent periods, we detect regional continuity of Early Neolithic lineages across Iberia, however the genetic contribution of hunter-gatherers is generally higher than in other parts of Europe and varies regionally. In contrast to ancient DNA findings from Central Europe, we do not observe a major turnover in the mtDNA record of the Iberian Late Chalcolithic and Early Bronze Age, suggesting that the population history of the Iberian Peninsula is distinct in character.

## Introduction

The changeover from a hunter-gatherer lifestyle to a productive mode of subsistence first emerged around 10,000 BCE in the Near East^1–3^. This so-called ‘Neolithic transition’ brought about fundamental changes in economy, social structure, demography and human health, and laid the foundations for agrarian societies and thus for ancient civilizations. Over the course of the 7^th^ and 6^th^ millennia BCE, agriculture spread from the Balkans to Central Europe. Another route of dissemination ran along the Mediterranean coastlines of Greece, Italy and the south of France to the Iberian Peninsula and north to the Paris Basin and Central Europe^4^. However, the process of Neolithisation was non-linear, with the archaeological record documenting influences of local cultural traditions^5^. In Iberia, the Neolithic transition, which began around 5700 BCE, appears to have been complex, and Mesolithic and farming communities coexisted and interacted for as long as two millennia^6^.

The artefactual remains, mainly ceramic, attest to the different origins and modes of Neolithisation on the Iberian Peninsula. On one hand stands a Mediterranean maritime colonization by Neolithic pioneers characterized by ceramic with clear parallels to the *Ligurian Impressa* collections of Italian origin^7–9^. Some Early Neolithic sites were also located in the hinterland, suggesting further routes of dissemination through the Pyrenees and/or along major rivers, such as the Ebro^10,11^. On the other hand, North African influences and contacts are tangible in the southern Iberian Neolithic^12,13^. All in all, Iberia appears a melting pot of influences and groups, combining Neolithic lifeways and with indigenous mechanisms of adaptation^14^.

In the Early Neolithic, we can observe common features shared over large areas, but also some regionally restricted phenomena^15^. In the whole territory, for example, there existed sophisticated systems of agriculture and livestock handling, with adaptable crops^16,17^ and seasonal strategies in flock management^18–20^. The groups of the *Franco-Iberian Cardial* and the *Epicardial* pottery styles appeared at this time and recent studies have revealed mutual diachronic influences in the material culture and economies of these cultures^15,21^.

An increasing number of burials from that epoch have come to light in the last years^22–24^. The oldest are individual inhumations, sometimes grouped in cemeteries^25^. It is also common to find human remains in caves, which were increasingly used for collective burials^26^. From the late 5^th^ millennium BCE onward, burial monuments appeared, and megalithic tombs became widespread^27^. This phenomenon links Iberia with other parts of Europe, indicating long-distance networks of communication. Meanwhile, certain parts of northeast Iberia maintained individual inhumations in pits, mostly in small cemeteries^28^. Besides megalithic tombs, ditched enclosures extending over more than 100 hectares started to dominate the landscapes of southern and central Iberia from 3300 to 3100 BCE, highlighting another widely spread European phenomenon^29,30^. During the Iberian Chalcolithic period (3000-2200 BCE), fortified settlements with stone walls and semi-circular bastions appeared in the western and southern parts of the Peninsula, while elsewhere, open settlements were still extant^31^. The diversity in settlement and burial types suggests the existence of social structures with different levels of complexity^32^. At the same time, as exchange networks, circulated precious goods such as ivory from Africa and even Asia to Iberia^33^.

From ca. 2600 BCE onwards, the so-called ‘Bell Beaker Phenomenon’ became manifest with its characteristic pottery, copper weapons, gold ornaments, and other prestige goods, an archaeological reflection of important social and economic changes which spread across vast regions of western and central Europe^34^. Iberia’s Bell Beaker assemblages are among the richest and most diverse in western Europe^35^, both in terms of settlements and burials^36^. It has thus long been a focus of archaeological research, commencing from migrationist hypotheses and leading up to current social explanations, where the Bell Beaker phenomenon is perceived as a package of prestigious objects exchanged and consumed by elite groups and displayed on special occasions^37^.

Around 2200 BCE, the Chalcolithic settlement and funerary practices were suddenly discontinued, particularly in the western and southern part of Iberia, where most of the ditched and fortified settlements were abandoned and collective sites and megalithic tombs were replaced by individual burials^31^. El Argar groups began to emerge in southeast Iberia, with large and massively fortified urban centers like La Bastida (Murcia, n. 28 on Fig. 1)^38^, which managed to control a territory of over 35,000 km^2^ during the following 650 years.

**Figure 1.**
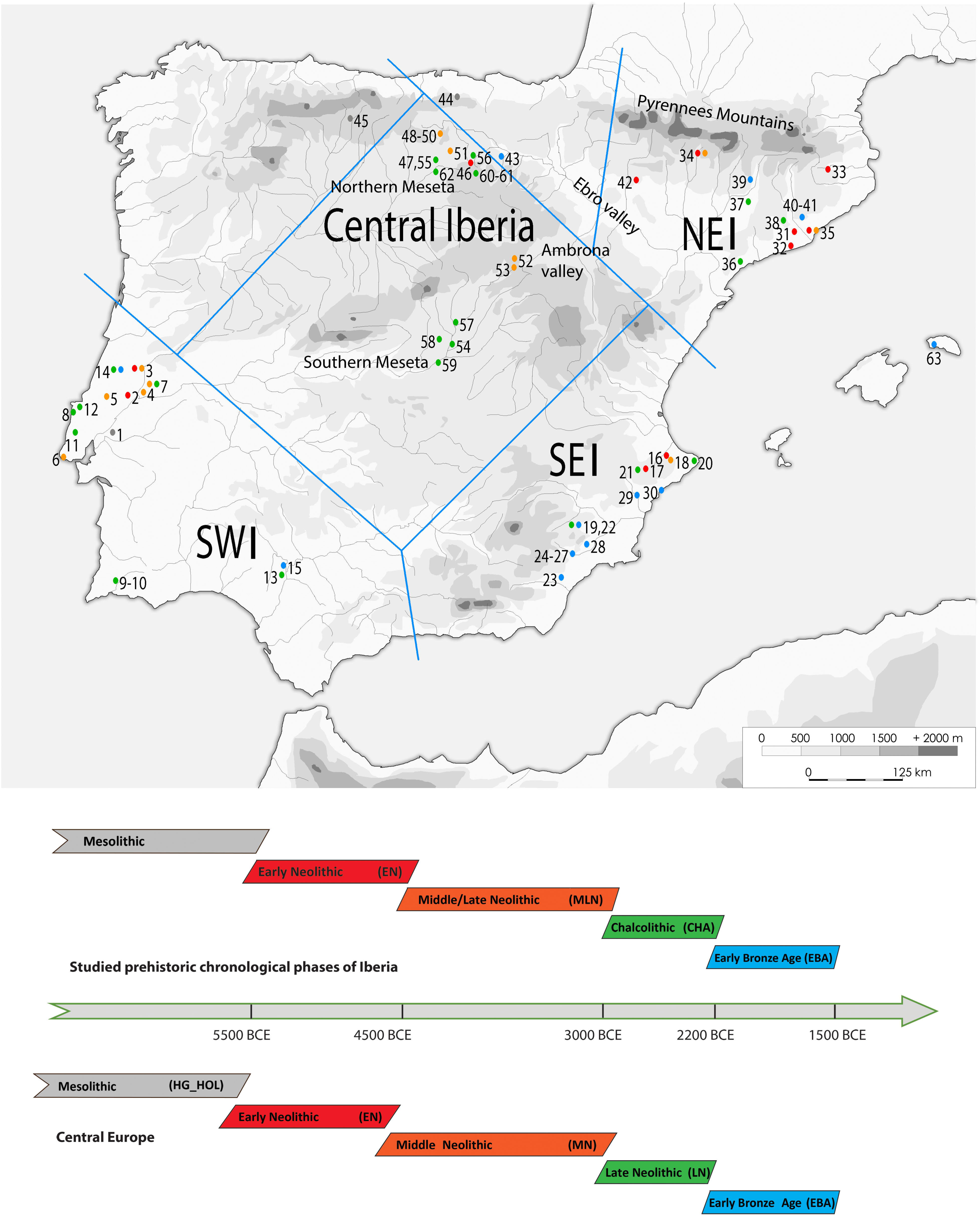
Map of the studied sites, including the published reference data and timing of archaeological periods on the Iberian Peninsula and in Central Europe. Geographic regions, also differentiated in the mtDNA analyses, are indicated as: NEI: northeast, SEI: southeast, SWI: southwest Iberia. Numbers on the map are colored according to the chronological periods, represented in the lower part of the figure. For the Central European chronology we used records from the most important comparative region of Central German Mittelelbe-Saale^55^. See Supplementary Table S5 for details. Site codes: 1. Moita do Sebastião, 2. Galeria da Cisterna (Almonda cave), 3. Gruta de Nossa Senhora das Lapas, 4. Gruta do Cadaval, 5. Gruta das Alcobertas, 6. Gruta do Poço Velho, 7. Gruta dos Ossos, 8. Tholos de Pai Mogo I, 9. Hipogeu de Monte Canelas I, 10. Hipogeu de Monte Canelas III, 11. Bolores, 12. Gruta de Malgasta, 13.Valencia Area 9, 14. Gruta do Carvalhal de Turquel, 15. Cobre las Cruces, 16. Cova de l`Or, 17. Cova de la Sarsa, 18. Cova d’en Pardo, 19. Molinos del Papel, 20. Cova del Barranc del Migdia, 21. Cova del Cantal, 22. Camino de Molino, 23. Fuente Alamo, 24. Lorca-Los Tintes, 25. Lorca-Madre Mercedarias, 26. Lorca-Castillo de Lorca, 27. Rincón de Moncada, 28. La Bastida, 29. Tabayá, 30. Illeta dels Banyets, 319. Cova Bonica, 32. Can Sadurní, 33. Cova d’Avellaner, 34. Els Trocs, 35. Sant Pau de Camp, 36. Barranc d’en Rifà, 37. Balma de Sargantana, 38. Cova de la Ventosa, 39. Cova de Montanissel, 40. Miguel Vives, 41. Can Gambús, 42. Chaves, 43. Valdescusa, 46. Alto de Rodilla, 47. Fuente Celada, 48. Fuente Pecina 1, 49. Fuente Pecina 2, 50. Fuente Pecina 4, 51. Alto de Reinoso, 52. La Mina, 53. La Tarayuela, 54. El Juncal, 55. Arroyal I, 56. El Hundido, 57. Camino de las Yeseras, 58. Humanejos, 59. Valle de las Higueras, 60-61. El Portalon, 62. El Mirador, 63. Es Forat de ses Aritges.

With regard to its population history, the Iberian Peninsula has been the focus of several recent archaeogenetic studies^39–44^. Studies focusing not only on mtDNA but also on Y chromosome markers have supported the model of a pioneer colonization in the north-eastern coastal regions of the Iberian Peninsula at the onset of the Iberian Neolithic^39,45^. On the other hand, the northern part of Spain (Cantabrian fringe) attested to a rather complex Neolithic transition^42^. Recent mtDNA and genome-wide analyses have put an emphasis on the genetic affinity and shared Near Eastern ancestry between the early Iberian farmers and the contemporary Central European Linearbandkeramik (LBK) population^46–48^. The latest mtDNA and genomic studies have revealed increased subsequent admixture of hunter-gatherer elements during the local middle Neolithic (La Mina; Alto de Reinoso, n. 51-52 on Fig. 1)^44,47^ and the Chalcolithic (El Portalon; El Mirador, n. 60-62. on Fig. 1) again reminiscent of processes observed in Central Europe^41,49,50^.

Despite numerous research projects being carried out over the past years, the Neolithic settlement history of Europe can still only be explained at a broad scale^47,49,51–54^. Regional transects though time detailing the developments and the course of the Neolithic in central Germany^55–57^ and in the Carpathian Basin^58^ have mostly been examined by mitochondrial DNA (mtDNA) control region data. Our project completes the latter series by focusing on the archaeological models and hypotheses that have been put forward for the Iberian Peninsula, and where diachronic (i.e. ‘through time’) sampling of ancient DNA allows the detection of demographic changes and discontinuities between 5700 and 1500 BCE. Essential research questions focus on three levels: i) the individual sites, ii) the Iberian Peninsula as a scene of Neolithic transition and iii) a comparison with contemporaneous ancient and modern-day Europeans. A key question of our study of the mtDNA diversity on the Iberian Peninsula through time was to examine to what extent regional and supra-regional cultural groups could be recognized as genetically identifiable entities, as shown in other areas of Europe^46,55^. A related question was whether the cultural breaks that can be seen, for instance, at the end of the Neolithic and Chalcolithic periods, were also accompanied by human population turnovers.

## Results

We processed ancient DNA samples of 318 human individuals from prehistoric Iberia and generated reproducible mitochondrial hypervariable region haplotypes (HVS-I, np 16020-16401) from 151 individuals following strict authentication criteria (see Methods). MtDNA haplogroup classification of further 62 samples was based on multiplex typing single nucleotide polymorphisms (SNPs) (Fig. 1, Supplementary Table S1-4). The DNA amplification and reproduction success of the HVS-I showed strong differences among samples from diverse regions of the Iberian Peninsula: the highest amplification success rates were observed in northeast Iberia (NEI) and higher altitude regions of central Iberia (CI) (71-78%), while especially the southern part of the Iberian Peninsula (southeastern Iberian (SEI) group and southwest Iberia (SWI) had very low amplification success rates (20-43%).

We merged these data with previously published HVS-I mtDNA results from 125 prehistoric individuals (Supplementary Table S5, S15), and separated the HVS-I dataset of 238, and mtDNA haplogroup information of 305 prehistoric Iberian individuals into geographically and chronologically defined groups (see Methods). The resulting haplogroup compositions of these groups are presented in Fig. 2.

**Figure 2.**
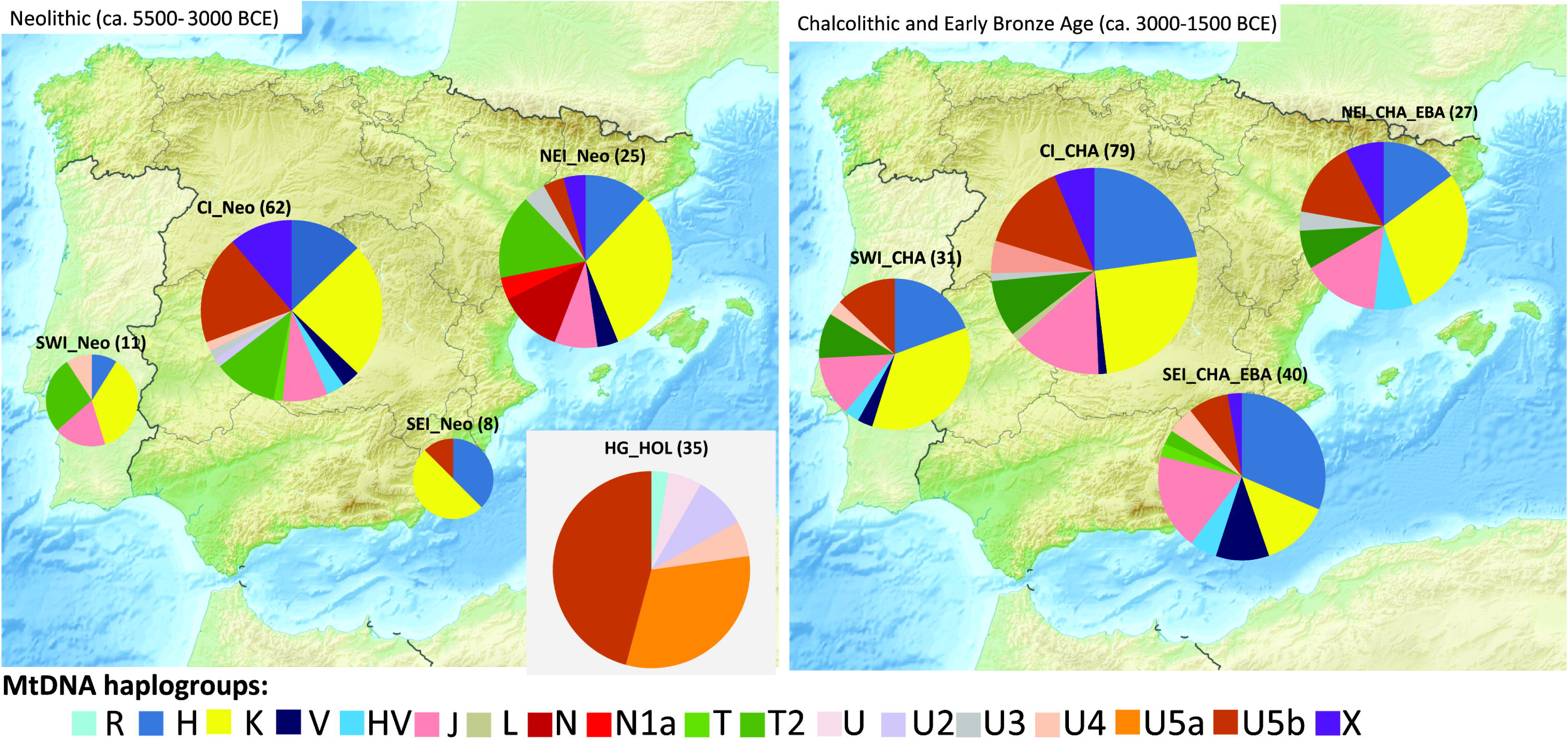
MtDNA haplogroup composition of the prehistoric Iberian groups. Abbreviations: Hunter-gatherers in Europe from the Holocene (HG_HOL), northeast Iberian Neolithic (NEI_Neo), northeast Iberian Chalcolithic and Early Bronze Age (NEI_CHA_EBA), central Iberian Neolithic (CI_Neo), central Iberian Chalcolithic (CI_CHA), southwest Iberian Neolithic (SWI_Neo), southwest Iberian Chalcolithic (SWI_CHA), southeast Iberian Neolithic (SEI_Neo), Chalcolithic and Bronze Age (SEI_ CHA_EBA). Relative haplogroup frequencies are presented in Supplementary Table S6. The background map of Iberian Peninsula was downloaded from Wikipedia (https://en.wikipedia.org/wiki/File:Relief_Map_of_Spain.png#file) and modified in Adobe Illustrator CS6 software.

### Haplogroup based analyses

The Iberian pre-Neolithic mtDNA substrate is still poorly understood. Recent NGS studies described hunter-gatherers belonging to haplogroup U5b in north Iberia^43,59^, while the suggested pre-farming presence of haplogroups H, N and U4^42^ will need to be verified by NGS-based whole mitogenome analyses. Therefore, we used Holocene period hunter-gatherers from Central, West and North Europe as proxy for the Iberian pre-Neolithic mtDNA substrate in our quantitative analyses (Fig. 2, Supplementary Table S6, S15). This continental hunter-gatherer dataset (HG_HOL, n=35) contains predominantly U haplogroups (U5b and U5a dominated, but also R, U2, U4, U* were observed^53,57,58,60–63^). In Iberia, haplogroup U5b was found most frequently in the CI Neolithic dataset (19.4%; n=62), and showed also relative high frequency in the CI Chalcolithic (19.9%; n=79) and NEI Chalcolithic-Early Bronze Age groups (14.8%; n=27).

The haplogroup composition of the Iberian Neolithic population shows similarities to the Early Neolithic data from Anatolia (6500-6000 BCE), the Carpathian Basin (5800-4900 BCE), and Central Europe (5500-4000 BCE, represented by the central German Mittelelbe-Saale region)^49,55,58^. Haplogroups K, J, T2, HV, V and X are observed in comparable frequencies in Iberian and Central European groups. However, the proportion of haplogroup H is higher in the Iberian Early Neolithic (EN) (22.2%; n=27) than in Central Europe (15%; n=160), while the frequency of N1a is very low (3.7% compared to 9.3% in Central Europe). Another difference with regards to Central Europe is the occurrence of haplogroup N* in Neolithic NEI (Supplementary Tables S6, S15).

We used the haplogroup frequencies of the studied groups for principal component analyses (PCA, see Methods). When compared to published ancient DNA data from the Near East, the western Anatolian Neolithic population is most similar to Central European Early-Middle Neolithic populations and to the Neolithic of northeast Iberia (Fig. 3, Supplementary Fig. S1-2), but it also shows some affinities with further Iberian farmers through common EN haplogroups (e.g. K, J, T2)^49^. The presence of haplogroup N* in the Pre-Pottery Neolithic in Syria and in NEI can also be interpreted in this light^39,64^, but this link needs to be verified by whole mitogenome analyses. Haplogroup compositions of all other Iberian groups are highly similar to each other, resulting in clusters on the PCA plots and Ward clustering tree. Our largest Iberian datasets are from Neolithic and Chalcolithic of CI (n=62 and 79), which show an elevated frequency of U5b (19.4 and 13.9%) compared to the NEI Neolithic. Interestingly, we also observe probable hunter-gatherer haplogroups U2 and U4, whereas the characteristic EN haplogroups N*, N1a are missing in the CI region. The southeastern Iberian dataset (n=48) contains eight Late Neolithic individuals with haplogroups K, H and U5b and 28 individuals from the Chalcolithic and 12 from the Early Bronze Age (EBA). Similarly to the CI groups it shows a mixture of haplogroups of the NEI group, but some of them (N*, N1a, U3) are missing in SEI.

**Figure 3.**
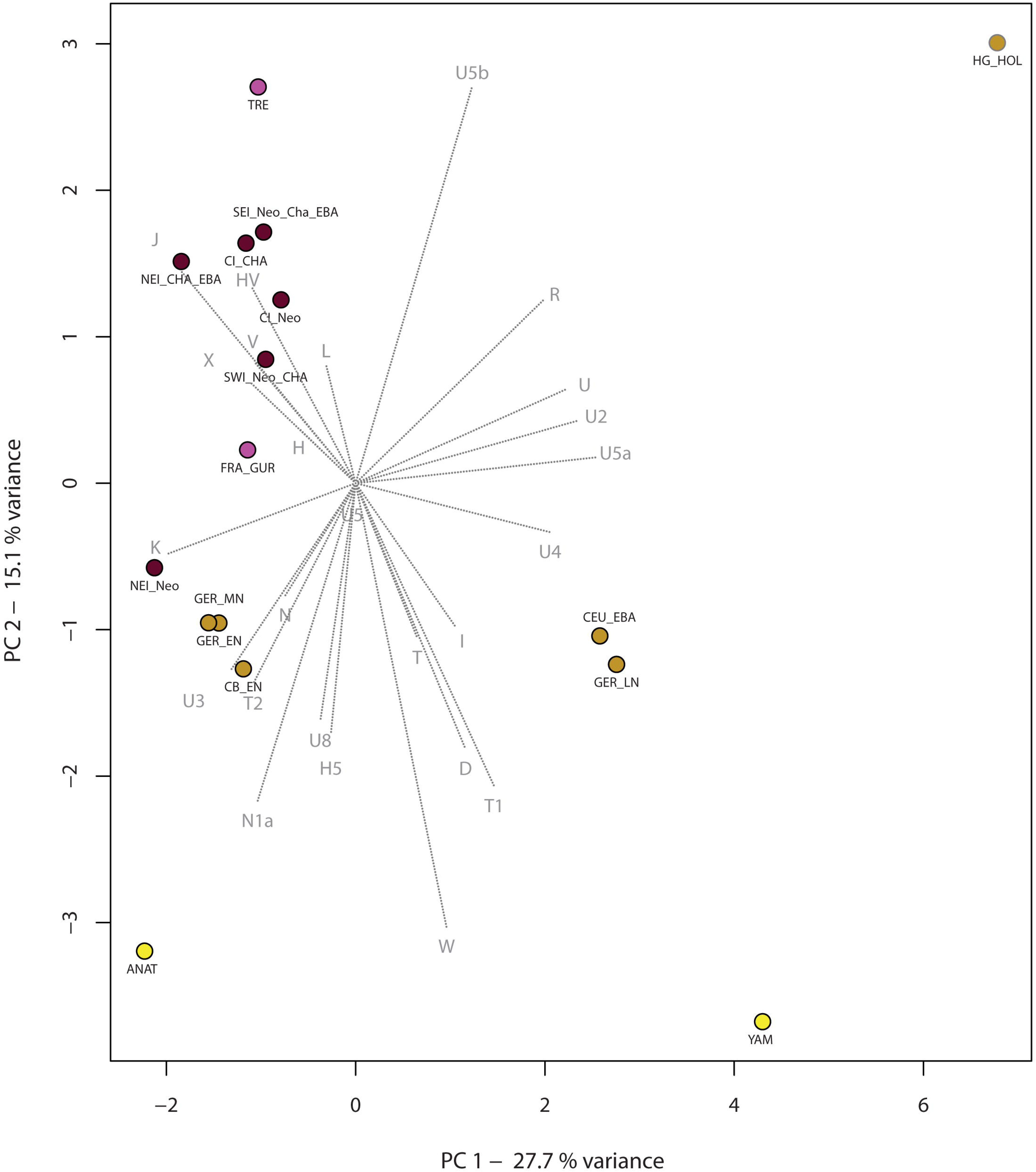
Principal component analysis based on haplogroup frequencies of 955 individuals from 16 prehistoric groups. The first two components display 36.6% of the total variance. Groups are colored according to their geographical positions: brown: Iberia, purple: France, ochre: Central and East-Central Europe, yellow: Eastern Europe and Near East. For abbreviations of Iberian groups, see legend of Figure 2. Further abbreviations: Central and North European hunter-gatherers from the Holocene (HG_HOL), Neolithic Anatolia (ANAT) Yamnaya (YAM), Early Neolithic Carpathian Basin (CB_EN), Early Neolithic Germany (GER_EN), Middle Neolithic Germany (GER_MN), Late Neolithic Germany (GER_LN), Early Bronze Age Central Europe (CEU_EBA), Neolithic Gurgy site in France (FRA_GUR), Neolithic Treilles culture in France (TRE). For further information see Supplementary Table S6.

The number of Neolithic and Chalcolithic SWI samples (n=12 and n=31) is too small to estimate the population dynamics in the region reliably (Supplementary Table S6). The combined dataset shows a predominance of haplogroup K (35.7%) and high frequency of haplogroups H, J and T (16.7-14.3-14.3%).

As Middle Neolithic (∼ 4500-3500 BCE) and Late Neolithic (∼ 3500-3000 BCE) periods of most sites are not separable at the current state of archaeological research, we combine these data in our analyses (MLN). Fisher’s exact (p=0.5473) and population continuity (p=0.3385-0.3722) tests confirm the connection between the EN and MLN periods of Iberia in general.

Some of the Iberian Neolithic mtDNA haplogroups (U2, N*, N1a) are not observed in the successive Chalcolithic Iberian population (n=156), whereas others maintain a steady frequency (V, T2, X) throughout 3500 years. An interesting exception is haplogroup L1b in the Late Chalcolithic Central Iberia at the site Camino de las Yeseras (n. 57 on Fig. 1), near Madrid. This group is most frequent in today’s West-Central Africa^65^, and hints at a connection to the North-West African coasts in prehistoric times.

In contrast to U5b, haplogroup U5a (common in Central European Mesolithic and Late Neolithic^55,57,66^) appears first in the 3000-2500 BCE period of central Iberia. Other typical Central European and Carpathian Basin Neolithic haplogroups such as H5, T1, W and U8, which are observed in later Central European Neolithic periods, are missing from the entire prehistoric Iberian dataset. These qualitative differences account for the separation of most of the Iberian prehistoric groups from the Central European Neolithic and Early Bronze Age populations along the second component of the PCA, shown on Figure 3.

Tests of population continuity and Fisher’s exact test between the whole Iberian Early and Late Chalcolithic datasets result in non-significant p values (p=0.5444-0.5578 and 0.9779), which support population continuity between the two periods. Unfortunately, a finer chronological separation of the Chalcolithic (CHA) dataset cannot be achieved for all samples and therefore we consider only a subset of the CHA dataset (n=71 for Early and n=47 for Late CHA) in this analysis (Supplementary Table S8).

We also tested whether this continuity could be the result of data merging across bigger geographic distance. However, the genetic continuity during the Early to Late Chalcolithic transition also holds at regional scale, as our population continuity test cannot reject continuity between the Neolithic (containing mainly Late Neolithic) and CHA groups of central Iberia (p=0.6033-0.6143), and NEI (p=0.9038-0.7835). This continuous genetic makeup is also supported by non-significant (p=0.2264 and 0.4484) differences in haplogroup compositions (Supplementary Table S7-8).

The Early Bronze Age sample set in the Iberian prehistoric transect is still very small (a total of n=37 individuals from all four regions studied). At the mtDNA haplogroup level the EBA does not show new influences or population changes at the onset of the Iberian Bronze Age according to results from Fisher’s test and test of population continuity, when both the entire Peninsula and southeastern Iberia in particular were considered (Supplementary Table S7-8).

A more detailed analysis of H sub-haplogroups focuses on 17 SNPs in the coding region of the human mitogenome. The largest proportion of the H individuals belongs to the subhaplogroup H1 (65.1%), and the second largest group is subhaplogroup H3 (14%), while 18.6% of the H individuals cannot be assigned to any of the subgroups included in the H-PLEX assay (see Methods, Supplementary Table S3-4)^67^. H3 was detected in Chalcolithic individuals from central, southeast and southwest Iberia. H1 was observed in each period and region, but more frequently in the Chalcolithic and Early Bronze Age than in the Neolithic. The comparative ancient H data from the European prehistory is too sparse for in-depth statistical analyses. However it becomes apparent from our results that the H diversity in prehistoric Iberia is different from the H diversity of Central Europe^55,58,68,69^, and more similar to the Neolithic populations in France^70,71^. Notably, common Central European subhaplogroups H5 and H7^68,69^ have not yet been observed in Southwest Europe.

Overall, we do not observe direct links between the Central European Late Neolithic (chronologically comparable to the Chalcolithic in Iberia) and the Iberian (entire and late) CHA groups at the haplogroup level, and thus cannot confirm a Late Chalcolithic expansion toward Central Europe as suggested by haplogroup H mitogenome data^68^.

### Haplotype and sequence based analyses

The mtDNA variation in prehistoric Iberia is further explored by HVS-I sequence and haplotype analyses of 238 individuals from the peninsula. The haplotype diversity of the hunter-gatherer populations of Central Europe (HG_PLEI (Pleistocene): Hd=0.879; n=29, HG_HOL (Holocene): Hd=0.931; n=35) provides the baseline for this comparative series. The diversity is lower in the Early Neolithic of Iberia (Hd=0.926; n=27) but increases in the following Middle-Late Neolithic (Hd=0.933; n=74) and Chalcolithic periods (Hd=0.944; n=118). The Early Bronze Age shows again a lower diversity (Hd=0.917; n=16), although this could be the result of the small number of individuals (Supplementary Table S9). The haplotype diversity among the regional groups is the lowest in the SWI Neolithic-Chalcolithic group in Iberia, and that along with the detected common basal J and K haplotypes could cause potential bias in lineage sharing results presented below.

The haplotype sharing (SHA) between the larger chronological groups detects no hunter-gatherer contribution in the EN group of Iberia, and a low level of hunter-gatherer maternal lineages in the MLN periods (6.8%) (Table 1). During the MLN, several new hunter-gatherer-type U5b lineages appear. Between the two successive Neolithic periods, the haplotype sharing amounts to 50%, and 43.2% of the MLN lineages are already present in the EN (as seen from our ancestral SHA). In the Chalcolithic period, we also observe a strong ancestral EN contribution (44.9%), with only 14.4% of the lineages originating from later Neolithic periods. The continuity of lineages is also seen in the smaller EBA dataset, which shows close connections to EN, MLN and CHA periods, and in which only 18.8% of the haplotypes are new (Table 1, Supplementary Table S10).

**Table 1.** Results of shared haplotype analysis (SHA): percentage of shared HVS-I haplotypes among the Iberian chronological groups (A), and ancestral haplotype analysis with the studied Iberian groups (B). Abbreviations: HG_HOL-hunter-gatherers, EN-Early Neolithic, MLN-Middle and Late Neolithic, CHA-Chalcolithic, EBA-Early Bronze Age, all in Iberia. For further details, see Supplementary Table 10 and Methods.

| **A: SHA** |  |  | **Detected in the following populations:** |  |  |  |  |
| --- | --- | --- | --- | --- | --- | --- | --- |
|  |  | **n sample** | **HG_HOL** | **EN** | **MLN** | **CHA** | **EBA** |
| **mtDNA lineages:** | **HG_HOL** | 35 | 100 | 0 | 6,756757 | 5,084746 | 0 |
|  | **EN** | 27 | 0 | 100 | 50 | 53,38983 | 28,125 |
|  | **MLN** | 74 | 31,42857 | 66,66667 | 100 | 52,54237 | 25 |
|  | **CHA** | 118 | 34,28571 | 66,66667 | 66,21622 | 100 | 40,625 |
|  | **EBA** | 16 | 0 | 33,33333 | 31,08108 | 43,22034 | 100 |
| **B: ancestral SHA** |  |  | **Detected in the following populations:** |  |  |  |  |
|  |  | **n sample** | **HG_HOL** | **EN** | **MLN** | **CHA** | **EBA** |
| **mtDNA lineages:** | **HG_HOL** | 35 | 100 |  |  |  |  |
|  | **EN** | 27 | 0 | 100 |  |  |  |
|  | **MLN** | 74 | 6,756757 | 43,24324 | 50 |  |  |
|  | **CHA** | 118 | 5,084746 | 44,91525 | 14,40678 | 35,59322 |  |
|  | **EBA** | 16 | 0 | 56,25 | 18,75 | 6,25 | 18,75 |

The Iberian early farming groups share generally high number of lineages (38.3-82.4%) with each other (Supplementary Fig. S3). The Central European Early Neolithic has the highest number of shared haplotypes comparing to the northeast Iberian groups, but the sharing is also high with prehistoric southeast Iberian groups. In contrast, lineage sharing with Holocene period hunter-gatherers is highest in the mostly Chalcolithic southeast Iberian group (8.6%) and in CI Neolithic (6.7%). Concerning temporal succession in NEI region, the proportion of hunter-gatherer lineages increases (from 0 to 4.8%) during the Chalcolithic period (Supplementary Table S10).

Continuity between Neolithic and Chalcolithic groups is reflected by the amount of shared lineages between the two periods in CI (61.2%), but both CI groups also share many lineages with all other Iberian groups (Supplementary Table S10).

Genetic distances (F_ST_) are generally low among the Iberian prehistoric groups, and none of them are significant (Supplementary Table S11). We observe the shortest distances from Central European EN-MN in northeast Iberia. By plotting the HVS-I based genetic distances after multidimensional scaling (MDS, Supplementary Fig. S4), large-scale relationships are displayed. Here, the NEI populations are within the cluster of the other Iberian groups, whereas the Central European and French datasets form a cluster that is slightly offset. Interestingly, MDS shows that some groups (e.g. NEI Neolithic) appear less differentiated from the remaining populations when compared to haplogroup-based PCA. The clustering of the groups on the MDS plot was therefore tested by the analysis of molecular variance. The best-supported groupings (i.e. with the highest variance among the clusters and lowest within the clusters) were evaluated by iteratively alternating the composition of the groupings. The Iberian groups differentiated from an EN Central European-Carpathian Basin and a MLN-EBA Central European-LN French clusters in this analysis as well, and can be further divided into a Neolithic-Chalcolithic CI and a NEI-SEI-SWI clusters (Supplementary Table S12).

We further compared HVS-I sequences of three larger chronological groups (EN, MLN and CHA) of the Iberian Peninsula with 133 modern populations. Genetic distances to modern-day populations are generally low, and restricted to certain region(s) of Europe (Supplementary Fig. S5, Supplementary Table S13). The Early Neolithic shows relatively high affinity to modern Afghanistan, Palestine, Iran, and Turkey (F_ST_ =0.00173-0.00428), but also high affinities to several European populations (*e.g.* Italian, French but also Belorussian). The genetic distances to modern populations increase with the MLN and CHA periods, but show general similarity to modern-day Europe. Recent genomic studies have highlighted the similarity of early farmers to modern South Europeans^72^, especially to modern day Sardinians^47,49^. This picture is not reflected in our mtDNA dataset, where Sardinians rank only the 67^th^ closest to the Iberian EN group out of 133 modern populations, and 46^th^ from the MLN group (Supplementary Table S13).

## Discussion

During the Last Glacial Maximum, the Iberian Peninsula, just as Southeast Europe and the Italian Peninsula, formed a classic Glacial Refuge Area for European populations, well documented by archaeological evidence^73–75^. Therefore, it is assumed that parts of today’s European population are the descendants of the residents of these refugia. This theory is based on coalescence date estimates for some mtDNA lineages (H1, H3, V, U5b1), which predate the Neolithic expansion, and also the high abundances of these lineages in Iberia today^76–80^. However, the Franco-Cantabrian glacial refuge theory has not yet been confirmed by ancient mtDNA or genomic analyses, and more Iberian Pleistocene and especially Holocene samples need to be investigated at full mitogenomic resolution.

With the end of the Last Ice Age and the beginning of the Holocene ∼12,000 BCE, a common European Mesolithic population emerged, which also included a distinct signal of Near Eastern origin^59^. The indigenous Iberian late Upper Paleolithic and Mesolithic populations were represented by whole mitogenome haplotypes assigned to haplogroup U5b from Northern Spain ^43,59^.

The Iberian Peninsula is characterized by diverse landscapes with distinct economic potentials. During the initial phase of Neolithisation, first farmers probably arrived in northeast Iberia primarily along the Mediterranean route and from there spread along the coastline and rivers (e.g., the Ebro Valley) into the hinterland^7–9,81^. Our genetic data support a substantial influx of Neolithic immigrants to northeastern Spain, where farmer lineages are most abundant and diverse, and on to other regions, like the SEI and CI. We found the persistence of ‘hunter-gatherer’ mtDNA haplogroups in the Neolithic to be strongest in Central Iberia. These genetic results match the archaeological data on the mode of Neolithisation described for these areas, so that the fully established Neolithic Iberian communities had distinct hunter-gatherer components^16,82,83^. The further the early farmers advanced into the inner/central and southern parts of the Peninsula, the higher the proportion of indigenous HG lineages. Geography appears to have been a decisive factor in the advance of Neolithic lifeways. Prehistoric central Iberia (southern and northern Meseta, Ambrona Valley), however, was never an isolated region surrounded by mountains, but rather an important hub for innovations and impulses coming into Iberia^84^. On a genetic level, Neolithic central Iberia showed corresponding mitogenomic connections to northeast and southeast Iberian regions (Supplementary Table S7).

The Pyrenees are also an area of special interest with regard to the route of immigration of the first farmers. This study includes ten samples from the cave site of Els Trocs, dated to the Early and Middle Neolithic^20^. Due to the considerable temporal differences between the occupations, the respective sample sets can be considered as genetically independent. The ^14^C data of the six individuals from the earliest phase cluster closely (Supplementary Information). Genome-wide analyses of five of these EN individuals revealed the group to be early Neolithic immigrants^47,49,50^.

Haplogroups N1a, J, T2, K and V appear, while the typical Iberian HG mtDNA haplogroup U5b is missing. Therefore, we suspect the presence of a (still) isolated EN group at Els Trocs. The isolated geographic position in the Pyrenees suggests immigration of this community from north of the Pyrenees (or even from Central Europe via the Rhone Valley) rather than from the west along the Mediterranean coastline. This alternative gateway to Spain has also been proposed based on archaeobotanical evidence^17,85^. The first and (to our knowledge) sole Iberian appearance of the mtDNA haplogroup N1a in an adult from Els Trocs, which matches identical HVS-I N1a haplotype in Anatolia and Central Europe, might have arrived on the continental route^47,49,55,86^. The shared roots between Southwest and Central Europe were especially conspicuous in the Northeast Iberian EN group, as the agreement in the common mtDNA haplogroups such as V, J, K, T2 and X suggests.

The transition from the EN to the MLN is documented in recent ancient genomic studies, which described an increase of hunter-gatherer elements in the farming populations of central Iberia during the MLN period at the site of La Mina and the Chalcolithic sites of El Mirador and El Portalón^41,47,49^. The Iberian Middle Neolithic genomic data are still too scarce (concentrated on a single site La Mina) to draw general conclusions about admixture with late hunter-gatherers^47^. La Mina people had 18.9-22.8% HG ancestry double than the EN individuals from Els Trocs (6.8-11%)^50^, and also higher than their contemporaries in Central Europe. These HG proportion further increased in the CHA to 27%^50^. In our study, we could support the general continuity of maternal lineages between MLN-CHA periods. The increase of HG maternal lineages through time is not seen on large scale SHA (Table 1), but observable in regional groups, such as in NEI, where the F_ST_ from HG_HOL decreases and the amount of HG_HOL haplotypes increases from the Neolithic to the CHA_EBA periods (0-4.76%). The same trend can be seen in our F_ST_ analyses where the CI_CHA (F_ST_ =0.17406) appear closer to the HG_HOL group than the CI_Neo group (F_ST_=0.21267) (Supplementary Table S7-8, S11).

According to the data currently available, the homogeneity of Chalcolithic ancient DNA results suggests that human mobility and genetic mixing had generally increased in Iberia by the Chalcolithic. Although we analyzed four geographically separated CHA groups in the northeast, central as well as in the southeast and southwest of the Iberian Peninsula, they did not exhibit any significant differences except for the comparison of the central and the southeastern group (Fisher’s test results in Supplementary Table S7, Fig. 2).

The detection of the ‘African’ Lb1 haplogroup at the Late Chalcolithic site Camino de las Yeseras (Madrid, central Iberia) is remarkable, given that ivory adornments of African origin have also been documented at this and other contemporaneous sites^87–89^. The observation of a western-central ‘African’ haplogroup alongside African artefacts and raw materials in Copper Age Iberia indicate long distance exchange that at least occasionally seem to have involved mobility of individuals and/or gene flow.

The Bell Beaker phenomenon was a decisive element in the Iberian Chalcolithic, lasting from the Late CHA to the EBA (2600 -1800 BCE)^35,36^. Our tests of population continuity and Fisher’s tests between Early and Late CHA periods supported population continuity between the two phases. However, our data structure did not allow studying the Early to Late CHA population changes at a regional scale, so that further studies might reveal local variation with regard to individuals with Bell Beaker / non-Bell Beaker cultural affiliations. Interestingly, we did not find evidence for direct genetic links between Chalcolithic Iberia and contemporaneous Central Europe. Links between the two regions had previously been suggested based in particular on Bell Beaker elements present across a wide geographic range^90–92^, as well as by the maternal genetic analyses of the El Mirador site^40^.

Interestingly, we also do not find evidence for influx in the East to West direction, as none of the investigated Chalcolithic individuals show ‘steppe ancestry’, which seen in contemporaneous Central European Corded Ware and Bell Beaker groups, suggesting that eastern influxes did not reach the Iberian Peninsula until later periods^49^. An evaluation of this unexpected observation will require further in-depth palaeogenetic studies.

Around 2200 BCE, the emergence of El Argar groups was evidently preceded by a break in Chalcolithic cultural traditions in southeast Iberia. Yet there are no apparent new influences or signals of substantial population change on the mtDNA haplogroup level at this time, so that the observed changes may either be due to an upheaval of existing social structures or an influx of groups that cannot be distinguished from the local population at the present level of genetic resolution, e.g., from southeastern Europe, as previously proposed for El Argar. Unraveling these apparently contradictory data will certainly require further in-depth analyses both on the archaeological and the archaeogenetic level.

## Conclusion

The present study, based on 213 new and 125 published mtDNA data of prehistoric Iberian individuals suggests a more complex mode of interaction between local hunter-gatherers and incoming early farmers during the Early and Middle Neolithic of the Iberian Peninsula, as compared to Central Europe. A characteristic of Iberian population dynamics is the proportion of autochthonous hunter-gatherer haplogroups, which increased in relation to the distance to the Mediterranean coast. In contrast, the early farmers in Central Europe showed comparatively little admixture of contemporaneous hunter-gatherer groups. Already during the first centuries of Neolithic transition in Iberia, we observe a mix of female DNA lineages of different origins. Earlier hunter-gatherer haplogroups were found together with a variety of new lineages, which ultimately derive from Near Eastern farming groups. On the other hand, some early Neolithic sites in northeast Iberia, especially the early group from the cave site of Els Trocs in the central Pyrenees, seem to exhibit affinities to Central European LBK communities. The diversity of female linages in the Iberian communities continued even during the Chalcolithic, when populations became more homogenous, indicating higher mobility and admixture across different geographic regions. Even though the sample size available for Early Bronze Age populations is still limited, especially with regards to El Argar groups, we observe no significant changes to the mitochondrial DNA pool until the end of our time transect (1500 BCE). The expansion of groups from the eastern steppe^47,93^, which profoundly impacted Late Neolithic and EBA groups of Central and North Europe, cannot (yet) be seen in the contemporaneous population substrate of the Iberian Peninsula at the present level of genetic resolution. This highlights the distinct character of the Neolithic transition both in the Iberian Peninsula and elsewhere and emphasizes the need for further in depth archaeogenetic studies for reconstructing the close reciprocal relationship of genetic and cultural processes on the population level.

## Materials

The studied sites are distributed across the Iberian Peninsula, with an emphasis on the archaeologically relevant regions on the Mediterranean coasts, in central and northern Spain and in southern Portugal (Fig. 1). We targeted representative sites from the Mesolithic, Neolithic, Chalcolithic and the Early Bronze Age to cover 4,000 years of prehistory on the Iberian Peninsula. Given that the southern European climate is usually unfavorable for DNA preservation, we implemented a flexible sampling scheme contingent on amplification successes, extending sampling from sites with good ancient DNA preservation and/or including additional sites to obtain sufficient data. Sample specific context information were supplied by our colleagues and project partners in Portugal and Spain or collected from previously published papers. All archaeological sites, relevant radiocarbon dates, and individuals incorporated in the present study are listed in Supplementary Table S1-2.

Altogether, 318 individuals from 57 archaeological sites in Spain, Portugal and Morocco were sampled and analyzed for this study. Whenever possible we preferred samples from recent excavations over those that had been held at museum collections for prolonged periods of time. Teeth and bone samples were taken under clean conditions in Iberian museums, at the Institute of Anthropology at Johannes Gutenberg University in Mainz, or directly on site in the case of La Mina, Arroyal, Els Trocs and in part at La Bastida. Mitochondrial profiles of 37 individuals from our project (Alto de Reinoso (n=27), La Mina (n=5) and Els Trocs (n=5) were previously published by our research team^44,47,49^. Here we also report additional samples from the sites of La Mina and Els Trocs.

The chronological classification of the burials was based on the archaeological data. In order to avoid possible problems due to terminological inconsistencies, we used temporal (from Mesolithic to Early Bronze Age; based on relative chronology and absolute dating) and geographic groupings to assess the population mitogenetic data from the Iberian Peninsula. We further distinguished groups based on contextual archaeological evidence, e. g. subsistence strategies, such as hunter-gatherers vs. early farmers. We targeted collecting two to three teeth from each skeleton and sampled bone only when teeth were not available.

## Methods

### Ancient DNA sample preparation

Standard sample preparation protocols were used during the ancient DNA work in the Institute of Anthropology in Mainz^55^. Samples were first UV irradiated for 30 min each side and then, the surface was removed by shot-blasting with aluminium-oxide-abrasive. Samples were then ground to fine powder using a mixer mill (Retsch). The milling process was controlled by hydroxylapatite blank controls, which were treated as samples in the subsequent DNA extraction and amplification steps. Series of samples were tested from each archaeological site, and the amplification success defined any further ancient DNA work. In general, A and B samples from different teeth or bone fragments per individual were analyzed independently. In the sole case is the site of Cova del Cantal, A and B samples, i.e. repeat extractions, derived from the same bone fragment.

### Ancient DNA extraction and amplification

Of the total 48 extraction batches, 25 were performed using the phenol-chloroform DNA extraction method^55^ and 23 using the silica based DNA extraction protocol^68^.

We combined different sets of primer pairs for the amplification of the HVS-I of the mitochondrial genome. Well-preserved samples were amplified in two segments (np 16046-16401), average preserved samples in four fragments (np 15997-16409) and samples with very fragmented DNA content were amplified with a combination of six overlapping primer pairs. All HVR I-II primers are listed in Table S11 in Alt et al.^44^, PCR, amplicon purification and sequencing protocols were the same as reported in Brandt et al.^55^. 40% of the PCR products were cloned using pUC18 vectors and E.coli cells^55,86^, and these clones were re-amplified and re-sequenced.

In order to get precise basal mitochondrial haplogroup determinations, we used the GenoCoRe22 multiplex PCR and ABI PRISM® SNaPshot® typing protocol^56^, and typed 22 coding region positions of the mitochondrial genome. Samples assigned to the haplogroup H, were further sub-typed used the H-PLEX17 focusing on 17 sub-haplogroup H defining single nucleotide polymorphisms (SNPs)^67^. Note that recent mitochondrial phylogeny updates have complicated the interpretation of typing results: the H1 defining variant G3010A is now also reported for subhaplogroups H30b, H65a, H79a, H105a (Phylotree Build 17).

Samples belonging to haplogroup U were also tested for six coding region SNPs via multiplex amplification and single base extension protocols, and further categorized into subhaplogroups U2, U3, U4, U5, U5a, U5b, and U8 (Supplementary Table S4). Haplogroup T were also further typed in the coding region for information on T1 or T2 sub-haplogroups (C12633A-T1 and A11812G-T2) and sequenced separately^69^. Conditions of the U and T–Plexes are as follows: PCRs were set up in a volume of 25 µI, containing 1 x PCR Gold buffer, 8 mM MgCl_2_, 0.02 µg BSA, 500 µM dNTPs, 0.02 µM of each primer, 2 U AmpliTaq Gold® DNA polymerase, and 2-3 µI DNA extract. Cycling conditions were 95°C for 10 minutes, and 35 cycles of 95°C for 30 seconds, 56°C for 30 seconds, and 65°C for 30 seconds, followed by a final adenylation step at 65°C for 10 minutes for the U-Plex and 32 cycles and an annealing temperature of 59°C for the H-PLEX17. Amplification success was checked via gel electrophoresis. Purification and SBE-reactions for the U-Plex was set up according to the GenoCoRe22 protocol.

Individuals with consistent HVS-I profiles from both samples were amplified over the whole HVS-I at least three times from at least two independent extracts, producing between 6-18 independent and overlapping amplicons (depending on the primer pairs used). PCR products that showed ambiguous nucleotide positions were cloned to monitor possible background contaminations and DNA damage. In addition, for those individuals of the sites Alto de Reinoso and La Mina that showed the same HVS-I haplotype, we also amplified and sequenced the HVS-II region via four overlapping primer pairs^44^.

### Authenticity and contamination

To monitor contamination, sample mix up and other potential errors we used the following authentication strategy: 1. All HVS-I haplotypes were replicated from at least two independent DNA extracts obtained from two skeletal elements from nearly all sampled individuals. The HVS-I fragment was amplified by numerous overlapping PCR amplicons. PCR products with ambiguous sequences were cloned, and 4-8 clones per PCR were sequenced. 2. Due to the applied primer-systems, amplification of the HVS-I enabled the identification of contiguous sequences with 6-10 SNP calls in the overlapping regions. 3. The final haplogroup call was based on several independent amplifications targeting different loci of the mitochondrial genome (HVS-I and coding region SNPs). In all reproducible and reported cases, the HVS-I sequences confirmed the haplogroup assignment by the GenoCoRe22 SNP typing (and vice versa), as did the results of the H-PLEX17, U-Plex and T-Plex assays. 4. All haplotypes could be placed in the mitochondrial phylogeny (www.phylotree.org). In all cases where ancient haplotypes differed from the current representative haplotypes on the mtDNA phylotree by private mutations, the latter were double-checked by several PCRs and by sequencing in combination with other SNPs. 5. Characteristic post-mortem DNA damage-derived substitutions (C>T, G>A) were observed, but these substitutions were neither reproducible nor consistent across sequence assemblies and could easily be distinguished from true SNPs. 6. Altogether 51 of the 2376 (2.15%) blank controls were contaminated during the course of the three-year investigation. No contamination could be detected in any of the multiplex PCRs. Of the 51 contaminated blanks, 0.96% (17 out of 1764) were PCR blanks, 6.4% (20 out of 311) hydroxyapatite milling blanks, and 4.7% (14 out of 301) extraction blanks. All contaminated blanks were sequenced and compared to haplotypes of samples analyzed along with these PCR extraction blanks. Fourteen individuals that were amplified from two extraction events had to be excluded from further analyses due to signs of potential contamination. 7. The HVS-I region was sequenced from all relevant co-workers (archaeologists, anthropologists and geneticists) (Supplementary Table S14) and compared with samples and blanks. None of the samples matched with the haplotype of the main processor of the samples (C.R.), although co-workers with common haplotypes (basal J, H-rCRS) cannot be ruled out as possible source of contamination. However, we find no signs of systematic contamination by these workers and consider it unlikely that samples that share these haplotypes were affected in various independent experiments whilst parallel samples were not.8. Ten samples were captured for whole mitogenomes^47^, and these results corresponds to the PCR and clone based sequence results (Supplementary Table S16).

### Population Genetic Analyses

### Reference populations and clustering of the Iberian data

We split pre-Neolithic samples of Western and Central Europe into two temporal groups: a Pleistocene group (HG_PLEI, before 10,000 BCE)^60,63^, a Holocene group (HG_HOL, after 10,000 BCE) from Central and North Europe^53,57,58,60–62^. The Iberian hunter-gatherer dataset (n=3) is too small to form a separate statistical group^43,59^. As comparative ancient populations we used Neolithic (5800-4900 BCE) datasets from the Carpathian Basin (CB_EN)^58,62^, Early (GER_EN, 5500-4000 BCE), Middle (GER_MN, 4000-2650 BCE) and Late Neolithic (GER_LN, 2800-2050 BCE) series from Central Germany^53,55,56,86,94,95^ and from Central European Early Bronze Age period (CEU_EBA, 2200-1550 BCE)^55,93^. From France, we included two Neolithic groups from the 4^th^ and 3^rd^ millennium BCE (FRA_GUR and TRE)^70,71^. From the Eastern European steppe, we included an mtDNA dataset associated with the Yamnaya culture (YAM, 3000-2500 BCE)^47,93,96^. Published HVS-I and haplogroup data are presented in Supplementary Table S15.

From the Iberian Peninsula we used all published mtDNA data that were secured by SNPs from the control region of the mtDNA^39,40,44,45,47–49,97^, and combined them with our results. The dataset from Cova de Montanissel was only used in haplogroup based tests, due to insufficiently reproduced HVS-I results^98^. We divided the ancient Iberian dataset into the following geographical groups: southwest Iberia (SWI), southeast Iberia (SEI), central Iberia (CI), northeast Iberia (NEI). For sufficiently large sample numbers, we further sub-divided these into the following chronological groups: Early Neolithic (EN, ∼ 5700-4500 BCE), Middle and Late Neolithic (MLN, ∼ 4500-3000 BCE), Early Chalcolithic (∼ 3000-2600 BCE), Late Chalcolithic (∼ 2600-2200 BCE), and Early Bronze Age (EBA, ∼ 2200-1500 BCE). Where the chronology could not be defined precisely, we classified the samples into Neolithic (Neo) and Chalcolithic (CHA) periods. Note the differences in chronological terminology between Central Europe and the Iberian prehistory, mostly due to the absence of a distinct ‘Chalcolithic’ period in the archaeological terminology used in Central Europe (Fig. 1).

### MtDNA haplogroup frequency based tests

We used principal component analysis (PCA) to visualize the relationships between relative haplogroup compositions of the Iberian and other prehistoric datasets. PCA has the benefit of reducing the complexity of the entire dataset while maintaining an accurate representation of its variability. PCA was performed with 16 ancient populations comparing haplogroup frequencies of 25 mtDNA haplogroups (see Supplementary Table S6), and calculated in R version 3.1.3^99^, using a customized *prcomp* function.

Hierarchical clustering was used to construct clusters of the predefined prehistoric datasets, based on similarities (distances) between their haplogroup compositions. This analysis helps interpreting the PCA plots in form of dendrograms. Clustering of haplogroup frequencies was performed using Ward type algorithm and Euclidean distance measurement method. The result was visualized as a dendogram with the *pvclust* package in R.2.13.1^100^. Cluster significance was evaluated by 10,000 bootstrap replicates. Significance of each cluster was given as an AU (Approximately Unbiased) p-value, in percentage.

Fisher’s exact test is a nonparametric test for statistical significance^101^. The null hypothesis for each Fisher’s exact test was that the Iberian groups belonged to the same population. The resulting p-value thus described the probability of obtaining the observed haplogroup compositions if both groups were part of the same metapopulation. Fisher’s exact test was performed with absolute frequencies of the haplogroups observed in the Iberian dataset. We used all haplogroup results per group in this analyses and tested a series of pairwise comparisons of Iberian chronological and geographic groups, using *fisher.test* function in R v.3.0.3^99^.

The test of population continuity explores whether genetic differences between two populations from two or more consecutive time periods can be adequately explained by genetic drift alone or whether a discontinuity between the two populations has to be assumed^55^. The tests were performed for pairs of chronological groups in specific regions (central Iberia, northeast Iberia). It was also performed for the entire Iberian time transect, using ‘metapopulations’ from consecutive pairs of chronological periods (HG_HOL, EN, MLN, CHA (early and late), EBA). Effective population sizes Ne=1000, 10000 were tested, calculating with differences between terminal dates of the chronological periods. Further parameters of the tests are presented in Supplementary Table S8.

### MtDNA HVS-I sequence based tests

We also characterized the Iberian populations with standard diversity indices that were calculated in DnaSP^102^. The HVS-I sequence range between np 16048-16410 was used for the calculations (Supplementary Table S9).

Shared haplotype analysis^103^ examines the occurrence of identical lineages in different populations. For the ancestral shared haplotype analysis (ancestral SHA), we took the relative chronology of the studied groups into account and tracked first appearance of any given lineage and their transmission (or presence/absence) in subsequent, i.e. later populations. SHA and ancestral SHA were performed as described in^58^. Levelplot of the percentage of shared lineages was visualized in R version 3.1.3, using level plot function. First, we considered the same groups that were counted in the F_ST_ analysis. In the next step, we assembled all Iberian regions, and we differentiated chronological groups of the Iberian prehistory (Supplementary Table S10). These groups were put into chronological order and ancestral haplotypes were determined. A major drawback of this process is the limited resolution of H-rCRS haplotypes.. On the other hand, available haplogroup definitions of H, V, HV and U haplotypes were used to separate these HVS-I sequences.

F_ST_ values (‘fixation indexes’) measure the amount of genetic structuring/differentiation in populations. With FST values we aimed to test whether pairs of populations were genetically distinct, and panmixia could be ruled out as basal assumption. FST values between pairs of prehistoric populations were calculated using reproduced HVS-I sequences (np 16056-16390) in Arlequin v. 3.5, applying a Tamura and Nei substitution model with a gamma parameter of 0.117^103^. FST p values were calculated based on 10000 permutations, and post hoc adjusted to correct for multiple comparison by the Benjamini and Hochberg method, using the function *p. adjust* in R. 3.0.3^104^.

Multidimensional scaling (MDS) was used to translate the distances/dissimilarities between multidimensional objects, represented by FST values, into spatial (Euclidean) distances and to visualize them in two-dimensional space. MDS was performed in R v. 3.1.3 using *metaMDS* function in vegan library^105^. Distance matrix was calculated based on Slatkin F_ST_ values, which FST values were calculated using HVS-I sequences (np 16056-16390) in Arlequin v. 3.5, applying a Tamura and Nei substitution model with a gamma parameter of 0.117^103^. We included 16 ancient groups in this analysis (Supplementary Table S11), but excluded the Holocene group, which had the greatest genetic distances from the other Central European and Iberian populations, and thus compressed the other parts of the MDS plot.

Analysis of molecular variance (AMOVA), based on HVS-I sequences (np 16056-16390), was performed between a subset of the Iberian and the Central European prehistoric groups used in the MDS (HG groups and YAM were left out). The “among clusters” and “within clusters” variance, F_CT_, F_SC_ and their p values were computed in Arlequin v. 3.5^103^. Geographic and chronological groups were arranged into different models, consisted of 2-5 clusters that were assumed as plausible from the MDS plot, and AMOVA was conducted for each arrangement (Supplementary Table S12).

We calculated FST values between modern and three prehistoric Iberian populations and mapped the genetic distances by interpolating between the geographic coordinates of modern populations. We calculated genetic distances (FST values) randomly choosing a maximum of 140 sequences per population (n = 17494 sequences altogether), in order to balance the differences in sample sizes, and calculated F_ST_ values between prehistoric Iberian (EN, MLN, CHA) and 133 present-day populations. The analysis was performed in Arlequin v. 3.5, using a uniform sequence length (np 16068–16365), and a Tamura & Nei substitution model with a gamma value of 0.177 (Supplementary Table S13)^103^. F_ST_ values were combined with longitudes and latitudes according to context information from the literature. The F_ST_ values and coordinates were interpolated with the Kriging method implemented in Arcmap ArcGIS version 10.3.

## Acknowledgement

Concerning research in the Alto Ribatejo, authors wish to thank to Fundação para a Ciência e Tecnologia for the support for research on the dawn of farming in the Tagus valley (project “Moving Tasks Accross Shapes” – PTDC/EPH-ARQ/4356/2014), as well as the Geosciences Centre of Coimbra University (strategic project UID/Multi/00073/2013).

## Financial statement

This study was funded by the German Research Foundation (Grant no. Al 287/14-1).

## Author contribution

K.W.A., R.R. and W.H. conceived the study. C.R. performed the ancient DNA work. A.S-N. analyzed the data. C. T-R., Í. G-M-d-L., H. A. M., P. B. R., R. B. B., R. d. B. B., A. M. H-C., R. F. F., C. A. F., J. J. E., L. R., C. O., M. I. F., I. S., O. V., R. M., V. L., J. S. D., J. A. L. P., C. R-d-T. M., M. S. H. P., F. J. J. M., J. L. M., A. A. F., K. T. L., A. M. S., M. M. R., L. M. O., V. M. D. S. G., C. C., A. W., J. R. B., A. M., J. P. M., M. H. O., J. C. M-G., J. C. P. E., R. C-A. B., T. T., E. C. B., J. L. C., A. C. A., C. L. v. L-V., C. B. B., P. R. M., A. P., J. I. R-G., M. A. E. B., R.P., E. M. H., E. M. I., J. V. d. M., R. M. G., V. M. C., O. L. J., and M.K. provided samples, anthropological and archaeological information. S. Z., C. K. and P. H. performed further bioarchaeological analyses. G. B., E. B., S. F., H. M., J. K., C. R. H. gave critical input to the version to be submitted. A.S-N., W. H., R.R., R.G-P., M.A.R-G., A. M. H-C., S. L. P. and K.W.A wrote the paper. All authors have read and commented the manuscript.

## Competing interest

The authors declare no competing interest.

## Data and materials availability

Consensus HVS-I DNA sequences were uploaded to NCBI GenBank under the accession numbers KY446554 - KY446703.

